# Temperature influences the excretion time and parasitic load of *Trypanosoma cruzi* in the *Triatoma infestans* vector

**DOI:** 10.1101/2023.11.08.566164

**Authors:** B. Álvarez-Duhart, S. Clavijo-Baquet, L. Valenzuela-Perez, J.D. Maya, M. Saavedra, S. Ortiz, C. Muñoz-San Martín, A. Bacigalupo, P. E. Cattan

**Affiliations:** Programa de Magíster en Ciencias Veterinarias y Pecuarias, Universidad de Chile, Santiago, Chile. Correo electrónico; Departamento de Ciencias Biológicas Animales, Facultad de Ciencias Veterinarias y Pecuarias, Universidad de Chile, Santa Rosa 11725, La Pintana, Santiago, Chile. Postal code: 8820808. Carrera de Medicina Veterinaria, Facultad de Medicina Veterinaria, Universidad San Sebastián, Santiago, Chile. Correo electrónico; Sección Etología, Facultad de Ciencias, Montevideo, Uruguay. Correo electrónico; Laboratorio de Biología Celular y Molecular, ICBM. Facultad de Medicina, Universidad de Chile, Santiago, Chile; Programa de Farmacología Molecular y Clínica, ICBM, Facultad de Medicina, Universidad de Chile, Santiago, Chile; Facultad de Ciencias Médicas, Escuela de Medicina Veterinaria, Universidad Bernardo O’Higgins, Santiago, Chile; School of Biodiversity, One Health and Veterinary Medicine, University of Glasgow, Glasgow, Scotland, United Kingdom. Postal code: G12 8QQ; ICBM. Instituto de Ciencias Biomédicas Facultad de Medicina - Universidad de Chile

**Keywords:** Climate change, transmission, kissing bug, *Trypanosoma cruzi*, temperature variation, parasitic load

## Abstract

*Trypanosoma cruzi* is a protozoan parasite transmitted by triatomine insect vectors, which expel their infectious dejections when they feed, causing Chagas disease in humans. The transmission and incidence of this vector-borne disease depend on the vital traits of its vectors, including *Triatoma infestans*, the main vector in Southern South America. Being an ectothermic species, its metabolism and its vital traits respond to temperature fluctuations. Here, we evaluated if changes in the average and variability of temperature expected with climate change modify: (i) the extrinsic incubation period (EIP) of *T. cruzi* within the vector *T. infestans*, (ii) its parasitic load, and (iii) the probability that its dejections were *T. cruzi*-positive. We acclimated triatomines infected with Dm28c *T. cruzi* strain to two constant and two variable temperature treatments and measured *T. cruzi* in their dejections by qPCR over a 42-day period. We observed that individuals in warm-temperature treatments showed lower EIP and higher parasitic load than cold-temperature treatments. Also, temperature variability can increase the parasitic load peak in cold-temperature treatments. Consequently, in a climate change scenario, there might be an increase in the vector capacity of *T. infestans* and probably a change in the risk of vectorial transmission of *T. cruzi*.

## Introduction

The relation between climate change and infectious diseases is one of most relevant ecological problems (Araya-Osses et al., 2020; Ayala et al., 2019; Lafferty, 2009). Vector-borne diseases are especially susceptible to temperature because their transmission rate and incidence are related to the biological traits of vectors, which are ectothermic arthropods (Lafferty, 2009). In this sense, climate change is expected to have an impact on multiple vector-borne diseases (Patz, 1996) such as Dengue fever (Struchiner et al., 2015), Chikungunya virus (Tjaden et al., 2017), Zika virus (Mordecai et al., 2019), West Nile virus (Paz, 2015), Bluetongue virus (Guis et al., 2012), Leishmaniasis (Chalghaf et al., 2018), Malaria (Kulkarni et al., 2016), and Chagas disease (Ayala et al., 2019). Temperature effect on the biting rate, reproduction, development, survival, and probability of becoming infectious after biting an infectious host (i.e., vector competence) has been reported on several disease vectors (Ciota et al., 2014; Damborsky et al., 2005; Luz et al., 1999; Shapiro et al., 2017). In summary, temperature shapes pathogen transmission, promoting its transmission at optimal constant temperatures and stopping it beyond lower and upper thermal activity limits (Lafferty, 2009; Stillwell & Fox, 2009).

Climate change projections include an increase in the average environmental temperature in the long term, and also more frequent periods of temperature variability, *e.g.* daily, monthly, or seasonal variation (Odoemene, 2017). Survival, reproduction, and thermal performance of ectotherms are affected by temperature variability in a non-linear way: temperature variation can positively affect individuals who are below their thermal optimum, and negatively affect individuals when they are above their optimum temperature (Clavijo-Baquet et al., 2021; Estay et al., 2014). Therefore, is complex to elucidate how climate change will modify the distribution and incidence of vector-borne diseases (Lafferty, 2009; Mills et al., 2010). Thus, to generate adequate predictions about it, the mechanisms that dictate the relation between temperature and vector traits must be fully understood (Mordecai et al., 2019).

The extrinsic incubation period (EIP, hereafter) is the time required for a pathogen or parasite to develop since its entry into the vector until the first forms capable of infecting another host appear (Tjaden et al., 2013). The EIP influences the incidence of vector-borne diseases (Christofferson & Mores, 2011; Mendes Luz et al., 2003; Rohani et al., 2009). This parameter determines the basic reproductive number R_0_ for vector-borne diseases (Hartemink et al., 2015). The EIP is also included in the vectorial capacity equation, which is a measure of the transmission potential of a vector-borne pathogen within a susceptible population (Christofferson & Mores, 2011; Dietz, 1993; Mendes Luz et al., 2003). Therefore, even small changes in EIP can have a large impact on the results of mechanistic models that are based on these equations (Hartemink et al., 2009; Pongsumpun, 2006; Putri et al., 2017). The practical relevance of this problem has been demonstrated for Dengue (Mendes Luz et al., 2003), Malaria (Paaijmans et al., 2010) and Bluetongue (Lambrechts et al., 2010), and the EIP of various species of parasites and viruses has been proven susceptible to temperature and its variability (Paaijmans et al., 2009; Rohani et al., 2009; Watts et al., 1987).

Chagas disease is a neglected tropical disease caused by the flagellate protozoan *Trypanosoma cruzi*, with 6 million people estimated to be infected in the world, predominantly in endemic areas of Latin America (Lidani et al., 2019; PHAO, 2022). This anthropozoonosis is transmitted primarily through the infected dejections of insects from the Triatominae subfamily (Hemiptera: Reduviidae) (Reisenman et al., 2010; Vieira et al., 2018). In the Southern part of South America, the most important vector is *Triatoma infestans*. Wild foci of *T. infestans* have been detected in Bolivia (Brenière et al., 2017), Argentina (Barbazan et al., 2010), Paraguay (Rolón et al., 2011), and Chile (Bacigalupo et al., 2010). Their presence poses difficulties for disease control efforts, due to their role in the recolonization process of peridomestic and domestic habitats (Antonella Bacigalupo et al., 2006; Bacigalupo et al., 2010; Brenière et al., 2016). In addition to the problem of wild foci, there is a projected rise in temperature due to climate change. In this sense, *T. infestans* temperature preferences range within 25-29 °C, (Lazzari, 1991). They display activity within a range of temperature between 18 and 42 °C (Canals et al., 1997).

Temperature is a major abiotic factor in *T. cruzi*-triatomine interactions (Asin & Catala, 1995). The development of *T. cruzi* and its relation with temperature has been studied under field and laboratory conditions (Asin & Catala, 1995; Giojalas et al., 1990; Tamayo et al., 2018), finding a positive relation between environmental temperature and the concentration of infectious forms in *Triatoma proctata* (Wood, 1954), *T. infestans* (Asin & Catala, 1995) and *Rhodnius prolixus* (Tamayo et al., 2018). An increase in temperature reduces the incubation time of the parasite within the vector (Asin & Catala, 1995). Field studies in Northwestern Argentina recorded higher numbers of infectious forms of *T. infestans* during the warm months (Giojalas et al., 1990).

However, the effect of temperature variability on parasite development has never been tested in controlled conditions for *T. infestans,* despite its relevance within the context of climate change. Currently, the minimum mean winter temperature in the Central region of Chile, where wild *T. infestans* foci have been reported, is 10.7 °C but it is also under strong daily temperature oscillation, with an average minimum of 4 °C to an average maximum of 16 °C for the coldest month (Araya-Osses et al., 2020). This species inhabits peridomestic habitats, which buffer temperature fluctuations, reaching up to ± 5 °C to ± 8 °C inside peridomestic structures and houses, respectively (Vazquez-Prokopec et al., 2002). Regarding climate change projections for the Central region of Chile, a temperature increase of 4 °C to 5 °C is projected in the worst-case scenario, while a conservative scenario predicts an increment of 2 °C (Araya-Osses et al., 2020). In this study, we evaluate how temperature and its variability influence the EIP and the parasitic load of *T. cruzi* in *T. infestans* dejections, and the probability of *T. cruzi*-positive dejections samples through time.

## Materials and Methods

A total of 144 *T. infestans* individuals in the fourth instar nymph were obtained from a colony reared at the Laboratorio de Ecología, Facultad de Ciencias Veterinarias y Pecuarias, Universidad de Chile. The fourth instar is easily manageable and has the capability to transmit *T. cruzi* to the vertebrate hosts (De Oliveira et al., 2018). The specimens were maintained individually in plastic flasks (3.8 cm x 6.8 cm) inside climatic chambers at 27 °C with a photoperiod of 12L:12D and 50 % to 70 % humidity. Prior to feeding on an infected mouse, the triatomines were fasted for 15 days.

### Triatomine infection with *Trypanosoma cruzi*

Triatomine infection was performed by means of blood ingestion from infected mice with *T. cruzi* clone Dm28c under controlled laboratory conditions. The trypomastigote forms of *T. cruzi* were cultivated in Vero cells (60-70% confluence) (ATCC® CCL-81TM), 5 ml of RPMI 1640 culture medium (Biological Industries) with 10% fetal bovine serum, plus 2 ml of trypsin, 1 ml of penicillin and 100 μg/mL of streptomycin (Biological Industries) as previously described (Valenzuela et al., 2018). Fourteen 8-week-old female BALB/c mice were inoculated only once, by intraperitoneal injection, with 1,000 parasites of the Dm28c clone (DTU TcI) in 100 ul/ml of RPMI 1640 medium. After inoculation, the mice were kept in cages at 20-25 °C with 40-70% relative humidity, with water and food *ad libitum* according to the bioethics protocol (N°: 19262-MED-UCH). Mice parasitemia was evaluated every 3 days from day 7 post-infection until reaching 80,000-500,000 trypomastigotes/ml to ensure triatomine infection. Once reached, the insects were fed on these *T. cruzi* inoculated mice. Balb/c mice were sedated following the bioethics protocol (N°: 19262- MED-UCH) dose of Ketamine 100 mg/ml (mouse dose 100 mg/kg) in association with Xylazine 5 mg/ml (mouse dose 10 mg/kg) and then put inside a plastic box (32 x 21 x 14 cm) over a heating pad. The triatomines were placed in the same plastic box with the mouse for 30 minutes in darkness, simulating nighttime. Each triatomine was weighed before and after the infection procedure to estimate the amount of ingested blood.

### Temperature treatments

After feeding on infected mice, all triatomines were randomly assigned to one of four temperature treatments (*i.e.* 18 °C ± 0 °C (n= 36); 18 °C ± 5 °C (n=36); 27 °C ± 0 °C (n=36); and 27 °C ± 5 °C (n=36)), set with an accuracy of ± 1 °C and a precision of 0.2 °C (PITEC^®^), with the same photoperiod and humidity as before. Temperature treatments were chosen considering the average temperature of the region (Araya-Osses et al., 2020) plus the temperature of the insect microsite habitat (Vazquez-Prokopec et al., 2002). In this sense, low temperatures in winter have been proposed as the most limiting factor in the population growth of *T. infestans* (De la Vega et al., 2015; Gorla, 2002; Gorla, 1992). Therefore, the lowest temperature to which we acclimated individuals resembled winter temperatures in their micro-environment (*e.g.*, a peri-domestic structure) adding 2 °C, which is equivalent to a conservative climate change scenario.

Individuals were acclimated to these low temperatures with and without the inclusion of temperature variability (*i.e.*, 18 °C ± 5 °C and 18 °C ± 0 °C, respectively). Additionally, since the thermal optimum for growth in *T. infestans* is easily reached inside of human houses (Canals et al., 1997), we also acclimated individuals to their optimal temperature, with and without temperature variability (*i.e.*, 27 °C ± 5 °C and 27 °C ± 0 °C, respectively). In the variable temperature treatments, variability was achieved during the light hours: the temperature started to increase linearly at 7:00 h, reaching its maximum at 8:00 h, then it was maintained constant, and began to decrease at 19:00 h, reaching its minimum at 20:00 h. The range of temperatures used in our experiments was chosen considering that the fluctuation range chosen lies well- within the species thermotolerance (Clavijo-Baquet et al., 2021). Temperature and humidity were monitored daily by the climatic chamber sensors.

### Dejection sampling

Samples of triatomine dejections were obtained from each flask every two days starting from 3rd day after infection until day 42. Dejection sampling was conducted mixing them with 80 µl of nuclease free water and storing them at -20 °C until DNA extraction. The triatomines were fed on non-infected laboratory mice (*Mus musculus*) at day 30 post-infection. We determine the refeeding day, considering the highly variable feeding frequency of triatomines in nature, ranging from months of fasting to daily intake (Cabello et al., 1988), and considering the effect of fasting on parasite survival (Kollien & Shaub 2000). After the experimental period, the insects were euthanized by freezing for a minimum of 48 hours following the bioethical protocol (N° 19262- MED-UCH).

### *Trypanosoma cruzi* detection in dejections

We extracted *T. cruzi* DNA from *T. infestans* dejections using the innuPREP Blood DNA Mini Kit (Analityk Jena AG), following the manufacturer’s instructions. All samples were co-extracted with 1 pg/μL of a 183 bp *Arabidopsis thaliana* (Brassicales: Brassicaceae) sequence generated by gBlocks® Gene Fragments (Integrated DNA Technologies IDT®) that was used as an exogenous internal amplification control, to evaluate carryover of inhibitors and DNA loss in the extraction process.

The qPCR assays amplified the satellite nuclear conserved region of *T. cruzi* with the primers Cruzi 1 (5 ’ASTCGGCTGATCGTTTTCGA 3’) and Cruzi 2 (5 ’AATTCCTCCAAGCAGCGGATA 3’) (Piron et al., 2007) in a final volume of 20 μL, containing 5 μL of template DNA, 5X HOT FIREPol® EvaGreen® qPCR Mix Plus (Solis BioDyne), 0.3 μM of each primer, and nuclease-free water. The cycling conditions were a pre-incubation of 15 min at 95 °C, followed by 40 cycles: a denaturation step of 95 °C for 15 s, a hybridization step of 60 °C for 20 s, and an extension step of 72 °C for 20 s, in a Rotor-Gene® Q (QIAGEN) thermal cycler. The emitted fluorescence was recorded at the end of each cycle, and a melting curve was run at the end of the program, with a ramp from 72 ° C to 95 ° C increasing 1 °C in each step, waiting for 90 seconds of pre-melting conditioning in the first step and 5 seconds for each subsequent step. Each reaction was carried out with a negative control (infection-free triatomine dejection’s DNA), a positive control (*T. cruzi* DNA quantification standard), and a no-template control (nuclease-free water). All samples were analyzed in duplicate.

The DNA standard curve for absolute quantification was obtained from the genomic DNA of *T. cruzi* strain DM28c. The calculation was estimated considering that a parasite cell contains approximately 200 fg of DNA (Kooy et al., 1989; Duffy et al., 2009). Serial 1/10 dilutions were made with nuclease-free water to cover a range between 10^6^ to 0.1 parasite equivalents/mL. All samples were co-extracted with 1 pg/μl of a sequence of 183 bp from tonoplast intrinsic protein 5;1 (TIP 5;1) of *A. thaliana* for the normalization of the parasitic load as previously described (Cabe et al., 2019). The quantification of the parasite equivalents from the DNA samples were calculated considering the amplification of the *T. cruzi* DNA standard curve and the results were normalized according to the results of the exogenous IAC. We chose this technique to detect *T. cruzi* infection because PCR-based methodologies are highly sensitive to monitor *T. cruzi* infection in triatomines in comparison to optical microscopy that has limited sensitivity in samples with low parasite numbers and present difficulties in differentiating *T. cruzi* from other morphologically similar trypanosomatids (Moreira et al., 2017).

### Statistical analysis

#### Extrinsic incubation period (EIP)

Only the first positive dejection sample for each individual was used to estimate the temperature effect on the EIP of *T. cruzi* in *T. infestans* (n=108). Generalized Linear Models (GLM) with a gamma function were fitted with EIP as a response variable, and temperature treatment (T), individual body mass before infection (m_b_), ingested blood during infection (b_i_) - estimated through the subtraction of the weight before and after feeding -, and mouse parasitemia (P_m_) as predictors. The goodness of fit of the models was evaluated by the likelihood ratio test (LRT) (Royle, 2013) and for model selection, we employed the Akaike information criterion (Akaike, 1973). Finally, to determine which treatments were different from each other, a *post hoc* analysis was performed using the Tukey test with Shaffer adjustment.

#### Parasitic load in *T. infestans* dejections

All positive dejections samples were included in this analysis (n=307). Generalized Additive Models (GAM) with a lognormal distribution were fitted for the parasitic load (Par-eq_ml_) as the response variable. In these models, temperature treatment (T), body mass before infection (m_b_), ingested blood during infection (b_i_), time in days of the collected sample post-infection (Day) and mouse parasitemia (P_m_) as predictors. Cubic splines were used to capture the nonlinear relationships between temperature treatment over time, ingested blood, and body mass before infection. We also included a random effect for triatomine individuals (ID) to correct the pseudo- replication of data. The model was selected according to the likelihood ratio test (LRT) (Royle, 2013) and Akaike information criterion (AIC) (Akaike, 1973).

#### Probability of positive dejections

We analyzed qPCR test results from all dejection samples (n=499) to determine their positive or negative *T. cruzi* qPCR test result, generating a binary variable. To test the probability of positive dejections over time according to the different temperature treatments of the individuals, we performed a Generalized Additive Model for Location, Scale, and Shape (GAMLSS) method for statistical analyses (R. A. Rigby et al., 2019; R. a Rigby & Stasinopoulos, 2008; Stasinopoulos & Rigby, 2007). This method does not require *a priori* assumptions about the shape of the relationship between the response variable and time (i.e., probability of positive dejections in time), fitting a cubic spline (cs). Thus, qPCR test results were the response variable; meanwhile temperature treatment (T), mouse parasitemia (P_m_), body mass (m_b_), ingested blood (b_i_), a cubic spline of the time (Day), and their interactions were the explanatory variables. We also included a random effect for individuals (ID) to correct the pseudo-replication of the data given by the effect of repeated measurements. In GAMLSS, the functions of the variables are unknown smooth functions, giving additional flexibility to the modeling process for a binary response variable. We performed models assuming a binomial distribution with ‘RS’ algorithm (R. A. Rigby et al., 2019). All analyses were performed in R version 2.13.0 (R Foundation for Statistical Computing, 2016) using RSTUDIO and the ggplot2, lmtest, multcomp, mgcv, pROC and gamlss packages.

### Ethics statement

All procedures with animals were conducted according to the Chilean animal protection law (Ley N° 20.380) and the Guide for the Care and Use of Laboratory Animals (Albus, 2012). The Scientific Ethical Committee for Research Safety of the Universidad de Chile approved all animal protocols used in the present work (Protocol N°: 19262-MED-UCH for Biosecurity and Ethical protocols).

## Results

### Extrinsic incubation period (EIP)

We found a significant effect of temperature over EIP (Tukey test; p < 0.05), but we did not find a temperature variability effect (Tukey test; p = 0.441) (Table 1, Fig. 1). The best model for EIP included the temperature treatment (T), the ingested blood during infection (b_i_) and the body mass before infection (m_b_) (Table S1, supplementary material). The predicted value for the EIP was 32 days for the cold constant treatment, 25 days for the cold variable treatment, 11 days for the warm constant treatment, and 13 days for the warm variable treatment (Table 1). An increase in ingested blood (b_i_) reduced the EIP, and its effect was greater for cold treatments (Fig. 2), decreasing the EIP to 29 days for individuals acclimated to cold constant treatment, and to 23 days for the cold variable one (Table 1).

**Figure 1.**
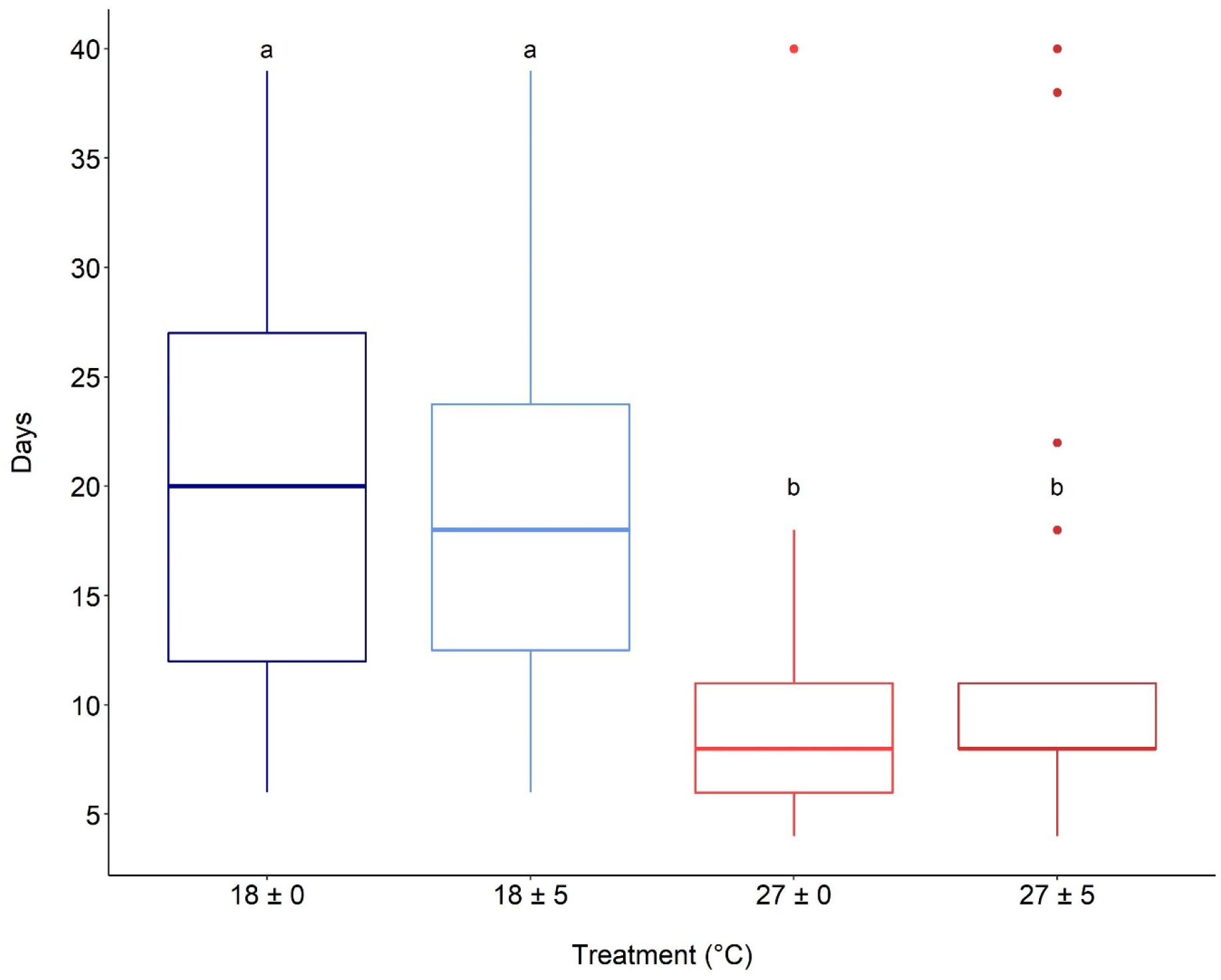
Boxplot of extrinsic incubation period (EIP) of *Trypanosoma cruzi* on *Triatoma infestans* exposed to four temperature treatments The letters show results from a posteriori test (Tukey test).

**Figure 2.**
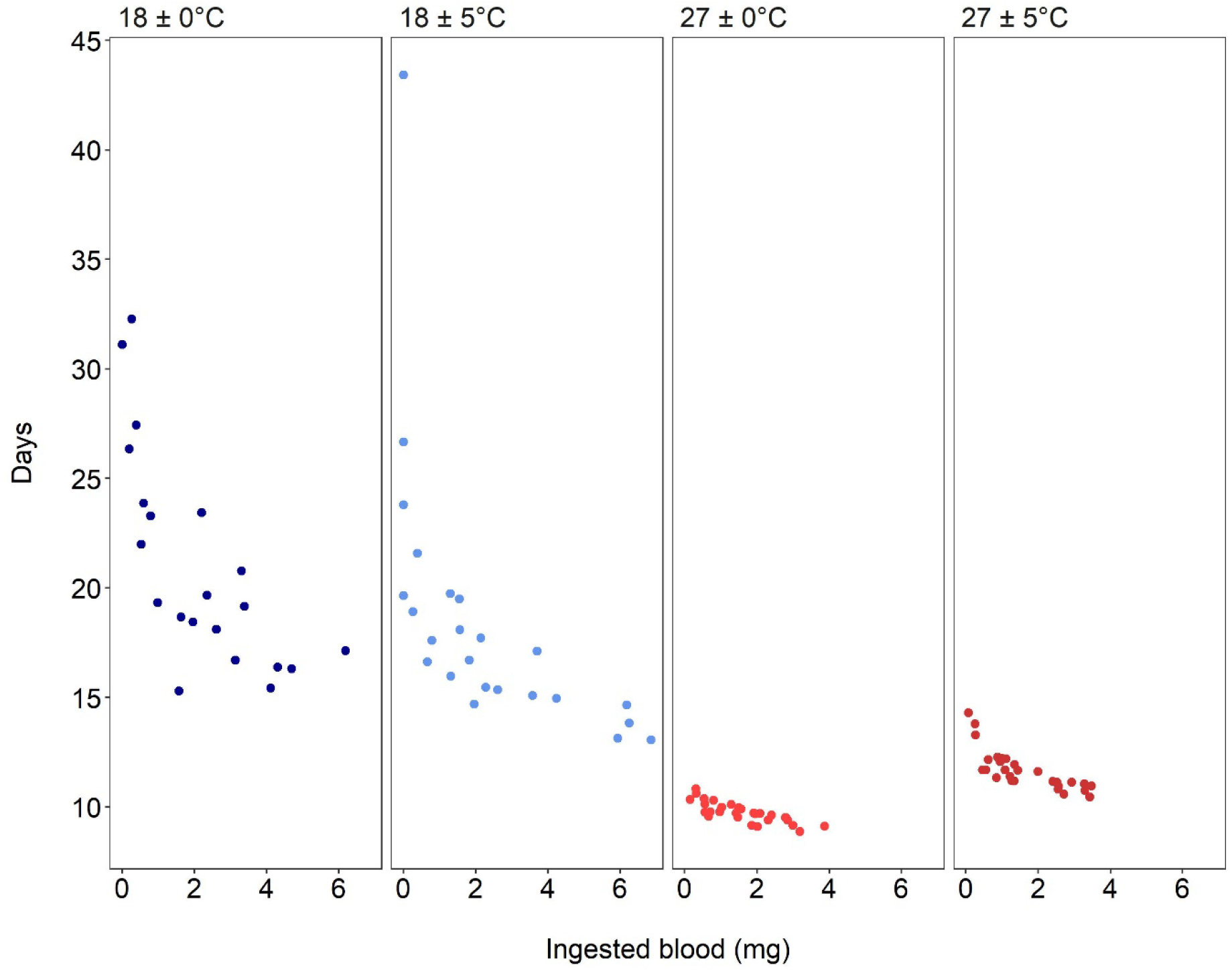
Extrinsic incubation period (EIP) of *T. cruzi* in *T. infestans* variation with ingested blood for four thermal treatments. Each treatment is illustrated on a different panel with a different color: cold constant temperature in blue (18 ± 0 °C), cold variable in light blue (18 ± 5 °C); warm constant temperature in red (27 ± 0 °C) and warm variable temperature in dark red (27 ± 5 °C). Ingested blood during the infection procedure (mg).

**Table 1.**
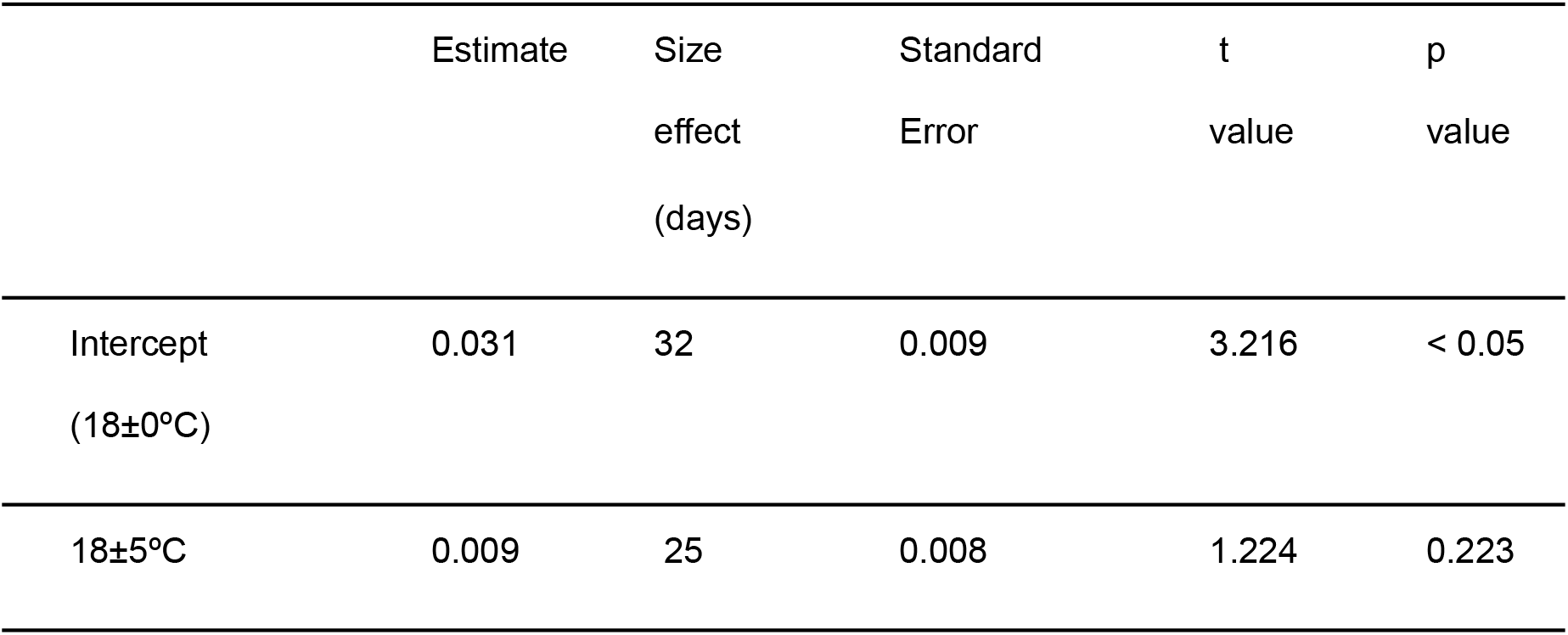
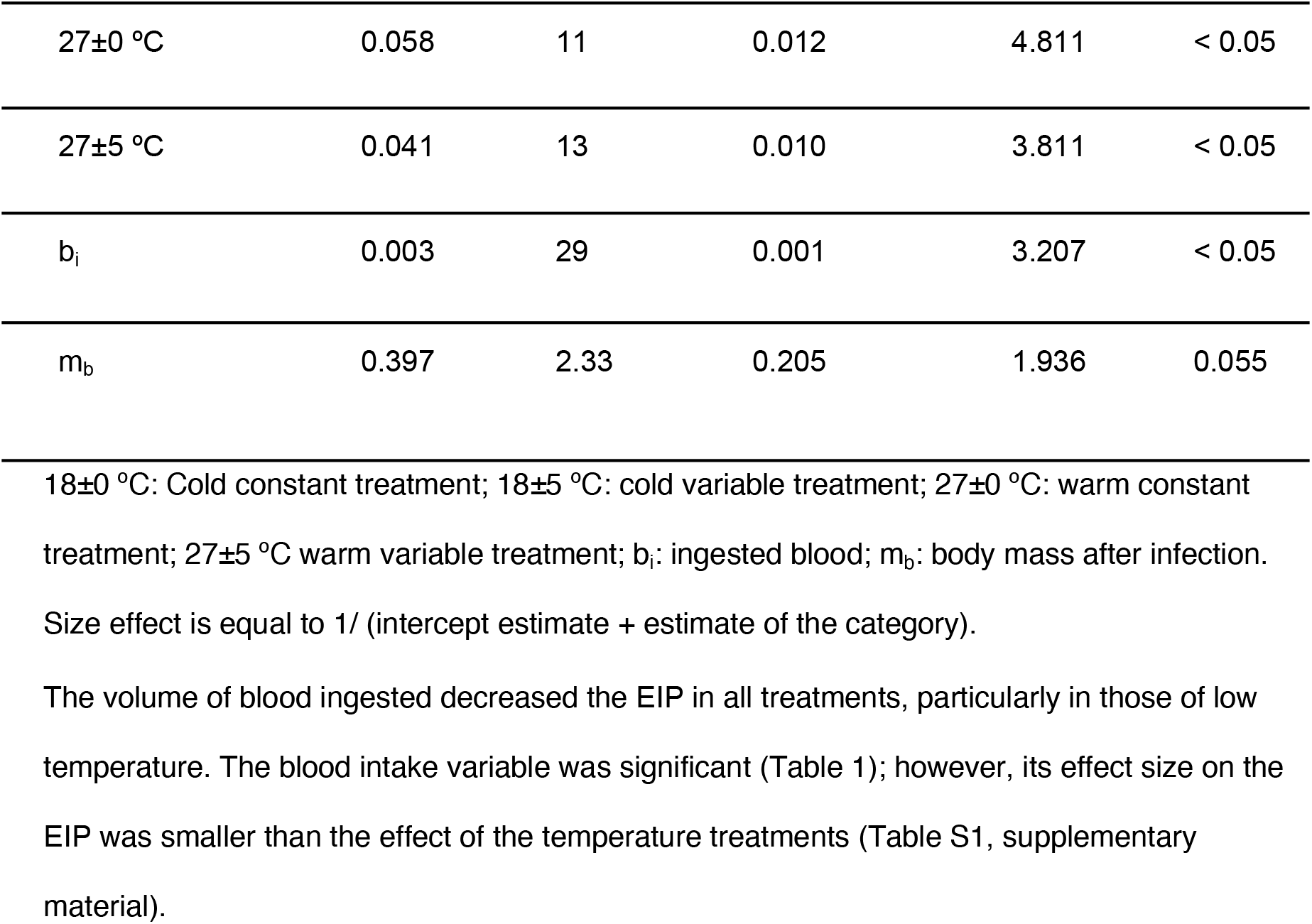
Summary of the best model fitted for *T. cruzi* EIP in *T. infestans*.

### Parasitic load in dejection samples of *T. infestans*

The parasitic load varied with time (Days) and temperature treatment (T) (Table S2 supplementary material). It describes a bell curve that starts low, peaks between 20 and 35 days, and then tapers off (Fig. 3). The curves change over time (days) according to the temperature treatment (Table 2, Fig. 3). Warm temperature treatments showed higher parasitic load and reached the peak of the curve earlier than the cold ones, being the cold constant treatment the one with lower parasitic load (Fig. 3 and Table 2). Nevertheless, temperature treatment effect over parasitic load was negligible (Table 2), and only significant along with time interaction (Fig. 3 and Table 2), meaning that bell-shaped curves for parasitic load were different across time (Fig. 3). Temperature variability showed an effect of increasing the parasitic load peak in cold treatments (Fig. 3). Treatments with temperature variability also showed a wigglier parasitic load curve, being more complex in the cold one (Fig. 3 and Table 2).

**Figure 3.**
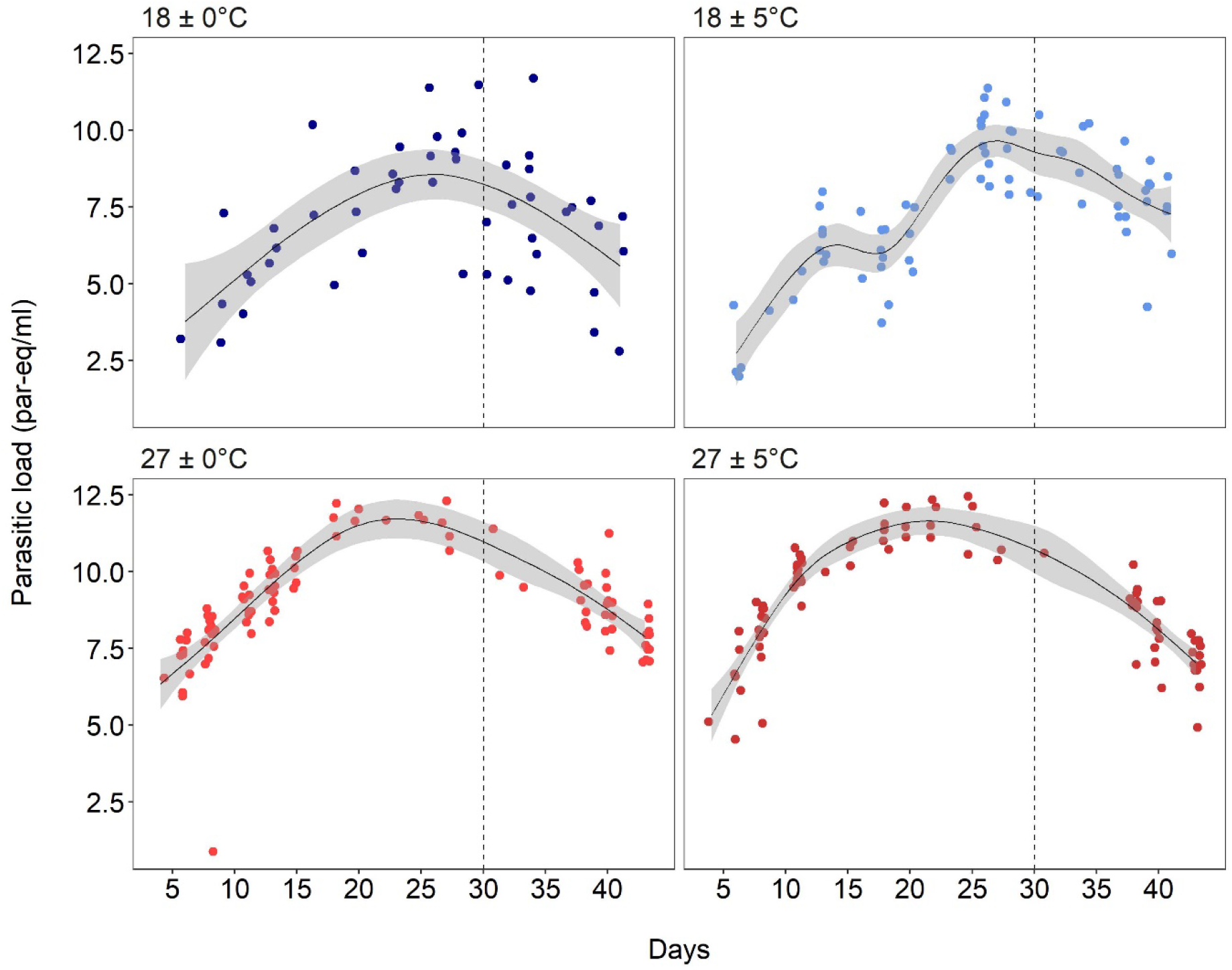
Parasitic load in *T. infestans* dejections samples throughout time (Days) by temperature treatment. Cold constant temperature in blue (18±0 °C), cold variable temperature in light blue (18±5 °C), warm constant temperature in red (27±0 °C), warm variable temperature in dark red (27±5 °C). The doted vertical lines mark the time of refeeding with an uninfected blood meal (day 30).

**Table 2.** GAM estimates and significance from the variables of the best model fitted for parasitic load of *T. cruzi* in *T. infestans*.

| Parametric coefficients: |  |  |  |  |
| --- | --- | --- | --- | --- |
| Variable | Estimate | Standard Error | t value | p value |
| Intercept (18±0 °C) | 5.1807 | 1.6984 | 3.050 | < 0.05 |
| 18±5 °C | 0.6251 | 2.2901 | 0.273 | 0.78510 |
| 27±0 °C | 2.9585 | 1.7775 | 1.664 | 0.09724 |
| 27±5 °C | 3.0884 | 1.7889 | 1.726 | 0.08547 |
| Approximate significance of smooth terms: |  |  |  |  |
|  | Edf | Ref.df | F | p value |
| s(ID) | 0.9506 | 1.000 | 5.392 | < 0.05 |
| s(Days): 18±0 °C | 2.8696 | 9.000 | 4.176 | < 0.05 |
| s(Days): 18±5 °C | 6.7396 | 9.000 | 8.433 | < 0.05 |
| s(Days): 27±0 °C | 2.9623 | 9.000 | 5.819 | < 0.05 |
| s(Days): 27±5 °C | 3.9339 | 9.000 | 6.957 | < 0.05 |
| s(m<sub>b</sub>) | 5.5756 | 6.694 | 3.793 | < 0.05 |
| s(b<sub>i</sub>): 18±0 °C | 3.1829 | 9.000 | 2.965 | < 0.05 |
| s(b<sub>i</sub>): 18±5 °C | 3.4857 | 9.000 | 1.208 | < 0.05 |
| s(b<sub>i</sub>): 27±0 °C | 4.9883 | 9.000 | 2.794 | < 0.05 |
| s(b<sub>i</sub>): 27±5 °C | 8.4865 | 9.000 | 1.935 | < 0.05 |
| R-sq.(adj) = | 0.568 | Deviance explained = | 63.4% | n =307 |

Estimate of the selected model; Standard error; t value; p value; Edf: effective degrees of freedom; Ref. df: Reference degrees of freedom; F: F-tests on smooth terms; p-value: p-value of the smooth terms; R-sq.(adj): Adjusted R squared value; 18±0 °C: Cold constant treatment; 18±5 °C: cold variable treatment; 27±0 °C: warm constant treatment; 27±5 °C warm variable treatment; s: spline; ID: triatomine individuals; Days: time measured in days; m<sub>b</sub>: body mass before infection; b<sub>i</sub>: ingested blood.

The best fit was achieved with the model including temperature treatment (T), Time (Days), body mass (m_b_), and ingested blood (b_i_) as predictor variables, with a spline in body mass (m_b_), time (Days), and ingested blood (b_i_), the last two per temperature treatment (supplementary material, Table S.3). The smooth term selected (Days, bs = "cs", by = T) implies that the parasitic load varied among temperature treatments (T) as time went by, and it was significant for all four treatments (Table 2). The ingested blood (b_i_) varied among temperature treatments (T) and all of them were significant (Table 2). The parasitic load was nonlinearly influenced by the body mass smooth term.

Altogether, according to the best-competing model (Table S2 supplementary material), the parasitic load trend changed with time and was affected by the amount of ingested blood, temperature treatment, and body mass (Table 2). The deviance of this model explained 63.4% of the variance, evidencing a good fit for the parasitic load in *T. infestans*.

### Probability of positive dejections

We obtained a total of 438 dejections samples from the 144 *T. infestans* individuals; 131 were negatives and 307 positives. For each treatment, the sample numbers were 78 for 18±0 °C (28 negatives and 50 positives), 117 for 18±5 °C (43 negatives and 74 positives), 117 for 27±0 °C (20 negatives and 97 positives) and 126 for 27±5 °C (40 negatives and 86 positives). We found a significant effect of temperature over the probability being positive, but we did not find an effect of temperature variability (Table 3, Fig. 4).

**Figure 4.**
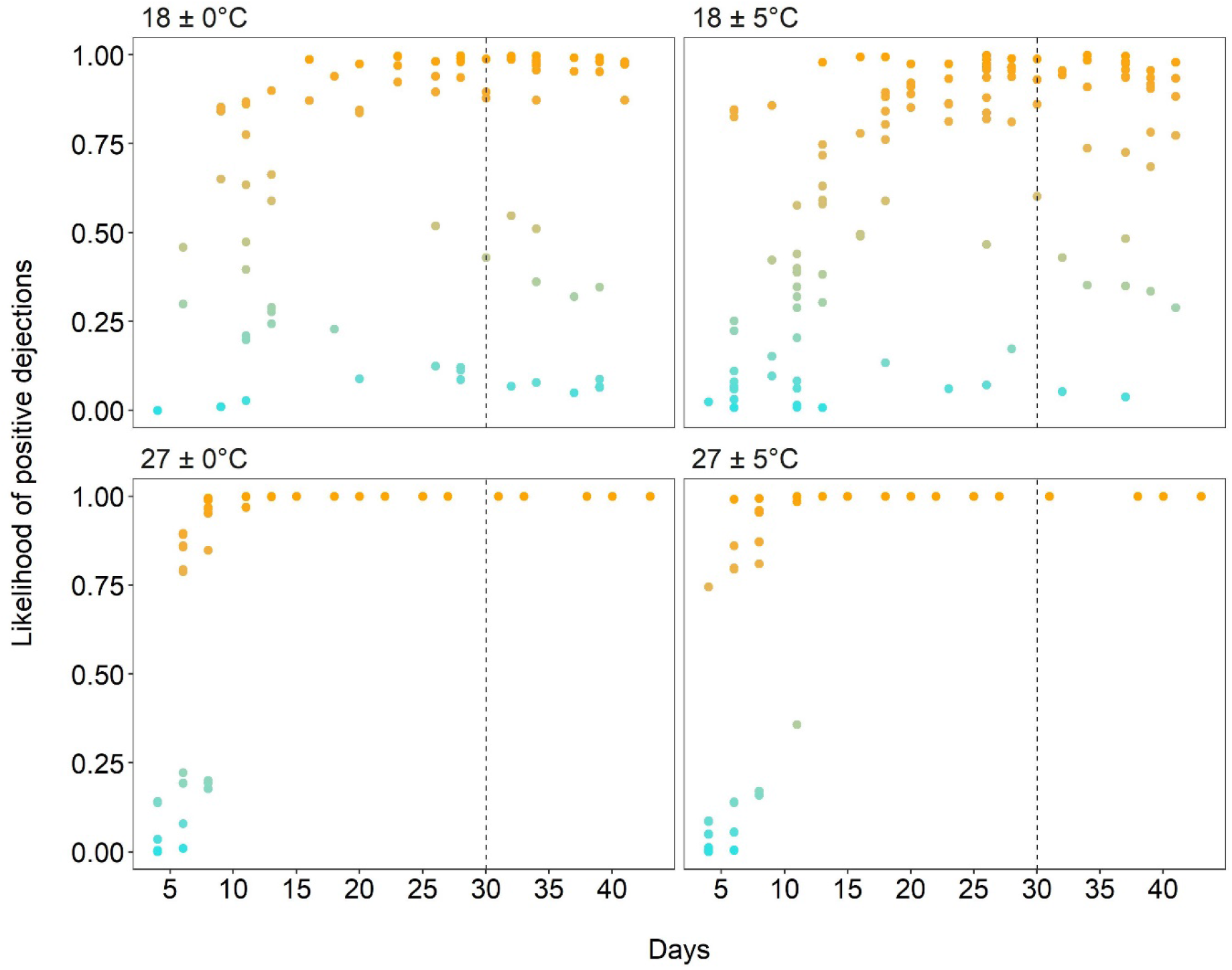
GAMLSS model. probability positive dejections to *T. cruzi* in *T. infestans* by temperature treatment. The probability is represented by a gradient palette from turquoise: minimum to orange: maximum probability of positivity. The doted vertical lines mark the time of refeeding with an uninfected blood meal (day 30).

**Table 3.** Parameters of the GAMLSS selected model.

|  | Estimate | Standard Error | t value | p value |
| --- | --- | --- | --- | --- |
| Intercept | -3.136 | 0.477 | -4038 | <0.05 |
| (18± 0 °C) |  |  |  |  |
| 18± 5 °C | 0.285 | 0.667 | 0.293 | 0.769 |
| 27± 0 °C | -5.474 | 2.573 | -2.317 | <0.05 |
| 27± 5 °C | -8.275 | 2.187 | -3.984 | <0.05 |
| cs(Days) | 0.151 | 0.020 | 4.682 | <0.05 |
| 18± 5°C:cs(Days) | -0.001 | 0.029 | -0.040 | 0.967 |
| 27± 0 °C:cs(Days) | 1.317 | 0.335 | 3.894 | <0.05 |
| 27± 5 °C:cs(Days) | 1.612 | 0.272 | 5.756 | <0.05 |

Estimate of the selected model; Standard error; t value; p value; 18±0 °C: Cold constant treatment; 18±5 °C: cold variable treatment; 27±0 °C: warm constant treatment; 27±5 °C warm variable treatment; cs: cubic spline; Days: time in days; ID: triatomine individuals

The best model fit for the probability of *T. cruzi* positive dejections included a cubic spline on time (Days) (i.e., cs (Days)) with an interaction between temperature treatments (T) (Table S3, supplementary material), showing differences between warm and cold treatments (p < 0.05) (Fig. 4 and Table 3). There was no significant difference between treatments with the same mean temperature on the probability of positive dejections (Table 3 and Fig. 3). As time (days) passed, the probability of positive samples exhibited an increase (Fig. 4), represented as the interaction between time and temperature (Table 3). In turn, for these warm treatments, the probability of obtaining negative samples decreased with time, becoming practically improbable after day 10. On the contrary, for individuals acclimated to low temperatures, the probability of finding negative samples for these treatments was maintained throughout the study period (Fig. 4).

## Discussion

Our results show that as the temperature increases, the time in which the first positive sample is detected decreases (Fig. 1). Therefore, temperature significantly changed the extrinsic incubation period (EIP) of *Trypanosoma cruzi* within *Triatoma infestans*, but temperature variability did not have a significant effect on it. Furthermore, the probability of positivity of the dejections changed with temperature but not with its variability (Fig. 4). The results also show different patterns in the parasite load over time between the different heat treatments, both constant and variable (Fig. 3).

The reduction in the EIP because of high environmental temperatures has been observed in three different species of triatomines (Asin & Catala, 1995; Catala et al., 1992; Giojalas et al., 1990; Tamayo et al., 2018; Wood, 1954) and is also supported by our results. *Trypanosoma cruzi* parasites reared *in vitro,* evidence that their number increased in direct relation to temperature (Elliot et al., 2015). Unfortunately, *in vitro* experiments do not include triatomine immune factors or their microbiota, among other factors, which affect the actual performance of *T. cruzi* within vectors in nature (Diaz et al., 2016; Duarte-Silva et al., 2020).

Regarding temperature variability findings on the EIP, populations are expected to evolve physiological adaptations to local climatic conditions in heterogeneous environments (Vázquez et al., 2017), exhibiting plastic strategies that allow them to survive a broad range of temperatures (Bozinovic et al., 2016), given that *T. infestans* and its parasites inhabit environments subjected to major temperature variability (Melo et al., 2020). *Triatoma infestans* can be found in the Chaco and other regions, where environmental temperature varies in a range that surpasses by more than the 10 °C those applied in our temperature variable treatments; therefore, both vector and parasite must adapt to these variations. It is possible that the absence of significant difference on the EIP of *T. cruzi* between constant and variable treatments reflects its adaptation to that natural environmental temperature variability. Thus, the absence of an effect of temperature variation on the EIP could be due to a wider thermal response curve of *T. infestans*, in accordance with the idea that vector and parasite populations in highly variable temperature environments have wider operative ranges compared to populations inhabiting stable microclimate environments (Mordecai et al., 2019). Moreover, *T. cruzi* is continuously exposed to temperature changes throughout its life cycle (Marliére et al., 2015), because the insect vector does not regulate its body temperature and ingests warm blood periodically, and due to the parasite alternating between insect vectors and a mammalian hosts (Magdaleno et al., 2009).

The EIP was also affected by the volume of blood ingested, showing a negative relationship (Fig. 2). Bed bugs typically consume a large amount of blood in a single meal, resulting in a high consumption of hemoglobin (Melo et al., 2020); a higher intake of this metalloprotein could explain the lower EIP, since it participates in the replication and survival of the parasite (Melo et al., 2020; Merli et al., 2016). On the other hand, it is likely that a greater volume of blood consumed during a feeding increases the probability of ingesting parasites, which would make the insect face a more potent infection.

Our study is the first evidence that parasitic load in *T. infestans* dejections varied over time, describing a bell-shaped curve, with lower findings of *T. cruzi* in the first and last days of the study time, with a peak on the third week. Previous work showed a positive linear relationship of parasitic load though time (Asin & Catala, 1995; Tamayo et al., 2018). However, those studies are not directly comparable, one of them they used optical microscopy (Asin and Catala 1998), which has been reported as less sensitive than qPCR (Kirchhoff et al., 1996), and the other worked with another vector species - *R. prolixus* -, with its samples obtained from tissue instead of dejections. Prolonged fasting has been shown to be an important factor in reducing parasitic loads in triatomines (Kollien & Schaub, 2000). In our experimental design we refed all individuals on day 30 with uninfected blood. However, the curve continued to decrease despite the refeeding. This suggests that there is another factor associated with the decrease in parasitic load over time. Remarkably, this bell-shaped curve occurred in all temperature treatments, but with different levels of parasitic load among them.

The treatments with warm temperature presented a higher parasitic load than the cold ones during the entire study period (Fig. 3). Within the cold treatments, the effect of in accordance with the idea that vector and parasite populations of highly variable temperature environments over parasite populations is evident, showing an increase in the parasitic load compared to the constant treatment. This difference has also been observed for the malaria vector *Anopheles stephensi* infected with *Plasmodium chabaudi*. In both vectors, the number of sporozoites per oocyst and the sporozoites found circulating in the hemocele were affected by temperature variability (Paaijmans et al. al., 2010).

It has been observed that the growth of epimastigotes is optimal at 28 °C and that greater development occurs as the temperature increases (Magdaleno et al., 2009). Therefore, the 5 °C rise in our experiment could have stimulated parasite development within the vector. Elevated temperature also affects the immune response of triatomines, causing a decrease in the activity of Prophenoloxidase (proPO) and the enzymatic cascade of phenoloxidase (PO), involved in the defense mechanism against pathogens (González-Rete et al. al., 2019; Melo et al., 2020) which in turn, could be influencing the degree of infection within the vector. Both factors could be playing a role in the earlier appearance of the parasitic load peack found in the warm variable treatment.

Vector parasitic load determines the modulation of the immune response and the degree of infection in mammalian hosts (Hofmeester et al., 2019); with greater parasitic load, greater the infection, tissue damage and mortality (Borges et al., 2013). Within vectors, it has been observed higher parasitic load increases the vectorial competence (Chacon et al 2022). Finally, we found a significant effect of temperature over the probability of obtaining positive samples, without a temperature variability effect, which shows a similar effect to what was found on the EIP. We were also able to observe that for insects maintained at low temperatures there was a greater heterogeneity in the probability of finding positive dejections, showing an individual effect. Moreover, it is important to consider that *T. infestans* is a domestic vector that can inhabit houses, where temperature fluctuates less than in peridomestic environments (Vazquez-Prokopec et al., 2002), thus the temperature variation effects will be more relevant to peridomestic environments and sylvatic foci (Bacigalupo et al., 2006; Bacigalupo et al., 2010; Saavedra et al., 2022).

In this study, we observed that different temperatures (warm and cold), along with their variability, lead to an increase in the parasitic load on *Triatoma infestans*, modifying its EIP and therefore its vectorial capacity (Christofferson & Mores, 2011; Dietz, 1993; Mendes Luz et al., 2003). In the context of climate change, this vector might transmit Chagas disease more effectively in the near future. Nevertheless, it is important to acknowledge that fluctuations in temperature can exert an influence on the survival and vital rates of insects, particularly when straying from their optimal thermal thresholds (Carrington et al., 2013; Clavijo-Baquet et al., 2021; González-Rete et al., 2019; Lambrechts et al., 2011). Thus, to obtain more accurate projections for this vector under climate change, more studies with other determining factors are need it. Nonetheless, we hope that these results contribute to a more complete understanding of the impacts of increased on temperature and it’s variability on this Chagas disease vector.

## Authors contributions

## Acknowledgments

Agencia Nacional de Investigación y Desarrollo (ANID) Fondecyt Regular 1180940; Fondecyt Iniciación 11160839. ANID - Programa Becas - Beca Doctorado Nacional - grant number 21171202; Doctorado Becas Chile 2019 - grant number 72200391; Fondecyt Regular 1210359.

## 8. Tables

**Table S1.** Model selection for GLM models of *Trypanosoma cruzi* EIP in *Triatoma infestans* acclimated to four temperature treatments that differed in temperature mean and temperature variability. The one in bold is the selected model.

| Models for EIP | df | LRT | AIC |
| --- | --- | --- | --- |
| $\sim T + P_m + b_i \times m_b$ | 9 | - | 709.19 |
| $\sim T + P_m + b_i + m_b$ | 8 | <0.05 | 711.38 |
| **$\sim T + b_i + m_b$** | **7** | **0.794** | **709.11** |
| $\sim T + b_i$ | 6 | <0.05 | 711.60 |
| $\sim T$ | 5 | <0.05 | 717.16 |

GLM: Generalized linear model; EIP: Extrinsic incubation period; df: degrees of freedom; LRT: likelihood ratio test; AIC: Akaike's information criterion; T: Temperature treatments; $P_m$ : mouse parasitemia; $b_i$ : ingested blood; $m_b$ : body mass after infection.

**Table S2.**
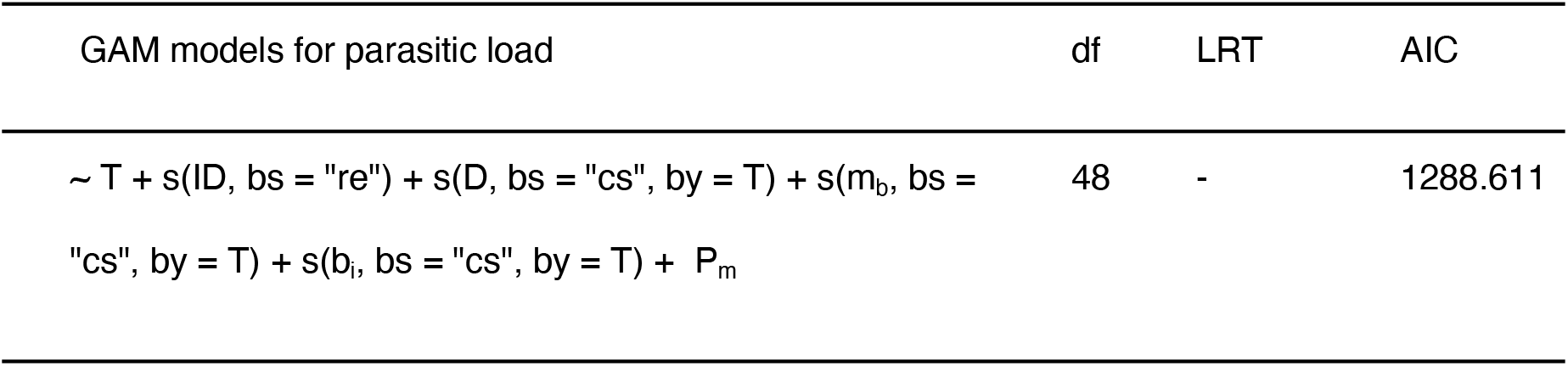
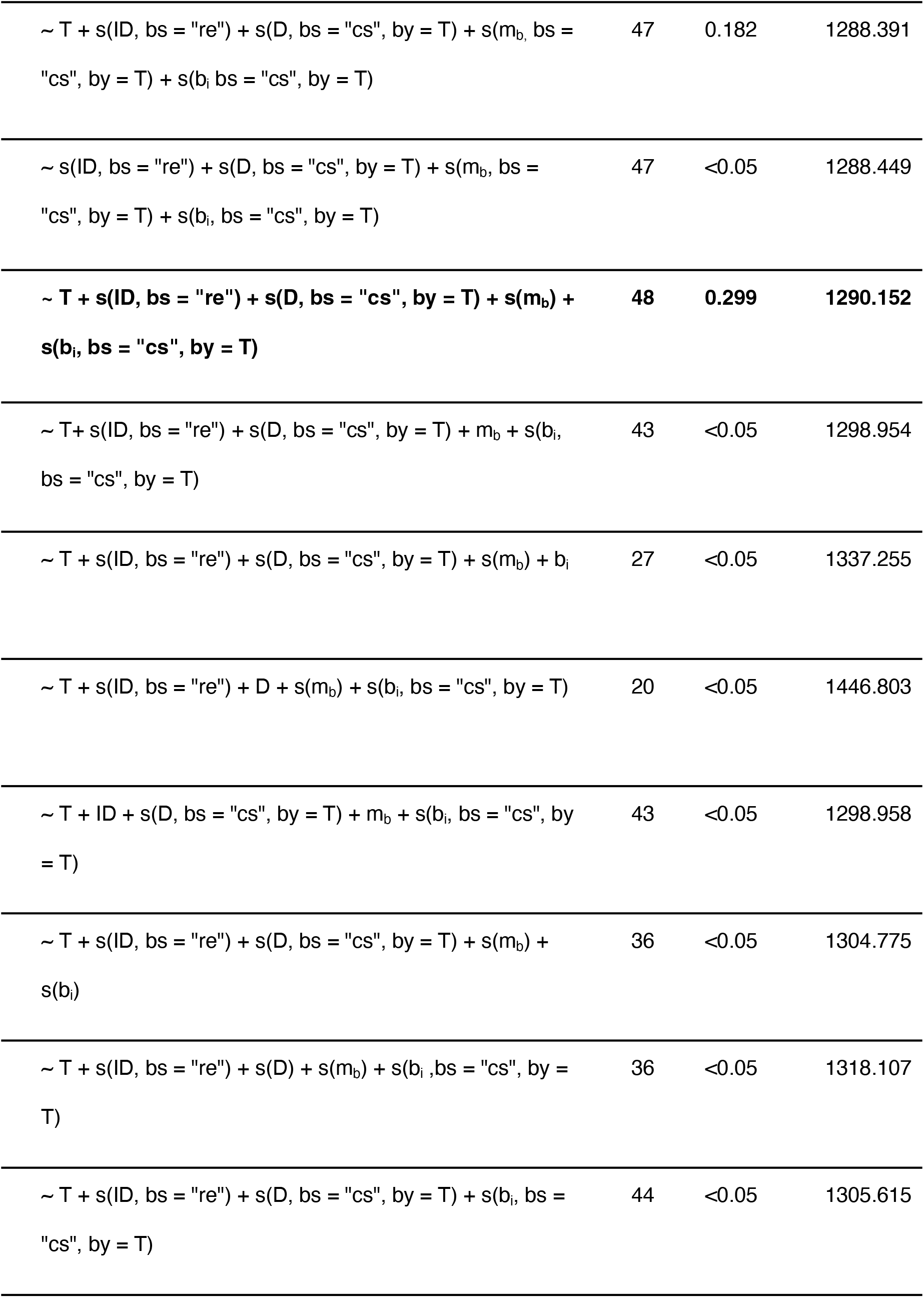
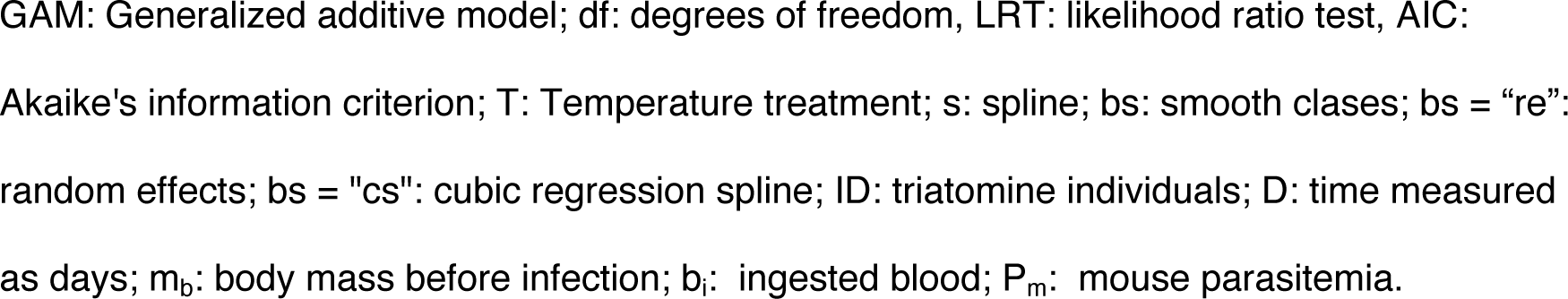
Model selection for GAM models of *T. cruzi* parasitic load in *Triatoma infestans* acclimated to four temperature treatments that differed in temperature mean and temperature variability. The one in bold is the selected model.

| GAM models for parasitic load | df | LRT | AIC |
| --- | --- | --- | --- |
| $\sim T + s(ID, bs = "re") + s(D, bs = "cs", by = T) + s(m_b, bs = "cs", by = T) + s(b_i, bs = "cs", by = T) + P_m$ | 48 | - | 1288.611 |
| ~ T + s(ID, bs = "re") + s(D, bs = "cs", by = T) + s(m<sub>b</sub>, bs = "cs", by = T) + s(b<sub>i</sub>, bs = "cs", by = T) | 47 | 0.182 | 1288.391 |
| ~ s(ID, bs = "re") + s(D, bs = "cs", by = T) + s(m<sub>b</sub>, bs = "cs", by = T) + s(b<sub>i</sub>, bs = "cs", by = T) | 47 | <0.05 | 1288.449 |
| **~ T + s(ID, bs = "re") + s(D, bs = "cs", by = T) + s(m<sub>b</sub>) + s(b<sub>i</sub>, bs = "cs", by = T)** | **48** | **0.299** | **1290.152** |
| ~ T + s(ID, bs = "re") + s(D, bs = "cs", by = T) + m<sub>b</sub> + s(b<sub>i</sub>, bs = "cs", by = T) | 43 | <0.05 | 1298.954 |
| ~ T + s(ID, bs = "re") + s(D, bs = "cs", by = T) + s(m<sub>b</sub>) + b<sub>i</sub> | 27 | <0.05 | 1337.255 |
| ~ T + s(ID, bs = "re") + D + s(m<sub>b</sub>) + s(b<sub>i</sub>, bs = "cs", by = T) | 20 | <0.05 | 1446.803 |
| ~ T + ID + s(D, bs = "cs", by = T) + m<sub>b</sub> + s(b<sub>i</sub>, bs = "cs", by = T) | 43 | <0.05 | 1298.958 |
| ~ T + s(ID, bs = "re") + s(D, bs = "cs", by = T) + s(m<sub>b</sub>) + s(b<sub>i</sub>) | 36 | <0.05 | 1304.775 |
| ~ T + s(ID, bs = "re") + s(D) + s(m<sub>b</sub>) + s(b<sub>i</sub>, bs = "cs", by = T) | 36 | <0.05 | 1318.107 |
| ~ T + s(ID, bs = "re") + s(D, bs = "cs", by = T) + s(b<sub>i</sub>, bs = "cs", by = T) | 44 | <0.05 | 1305.615 |

GAM: Generalized additive model; df: degrees of freedom, LRT: likelihood ratio test, AIC: Akaike's information criterion; T: Temperature treatment; s: spline; bs: smooth classes; bs = "re": random effects; bs = "cs": cubic regression spline; ID: triatomine individuals; D: time measured as days; $m_b$ : body mass before infection; $b_i$ : ingested blood; $P_m$ : mouse parasitemia.

**Table S3.** Model selection for GAMLSS models for the probability of *Trypanosoma cruzi* positive dejections in *Triatoma infestans* acclimated to four temperature treatments that differed in temperature mean and temperature variability The one in bold is shows the selected model.

| GAMLSS models for positive dejections | gl | AUC | AIC |
| --- | --- | --- | --- |
| $\sim \text{TxcS}(\text{D}) + \text{random}(\text{factor}(\text{ID})) \times P_m + \text{cs}(b_i) \times m_b$ | 102 | 0.9885 | 331.528 |
| $\sim \text{TxcS}(\text{D}) + \text{random}(\text{factor}(\text{ID})) \times P_m + b_i \times m_b$ | 100 | 0.9887 | 325.894 |
| $\sim \text{TxcS}(\text{D}) + \text{random}(\text{factor}(\text{ID})) \times P_m + b_i + m_b$ | 99 | 0.9887 | 323.865 |
| $\sim \text{TRTxcS}(\text{D}) + \text{random}(\text{factor}(\text{ID})) + P_m + b_i + m_b$ | 99 | 0.9887 | 323.865 |
| $\sim \text{TxcS}(\text{D}) + \text{random}(\text{factor}(\text{ID})) + P_m + b_i$ | 98 | 0.9887 | 322.044 |
| $\sim \text{TxcS}(\text{D}) + \text{random}(\text{factor}(\text{ID})) + P_m$ | 97 | 0.9888 | 320.207 |
| **$\sim \text{TxcS}(\text{D}) + \text{random}(\text{factor}(\text{ID}))$** | **96** | **0.9889** | **318.761** |

GAMLSS: Generalized additive model for location, scale and shape; df: degrees of freedom, AUC: Area under the ROC Curve, AIC: Akaike's criterion; T: Temperature treatment; cs: cubic spline; D: time measured as days; ID: triatomine individuals; $P_m$ : mouse parasitemia; $b_i$ : ingested blood; $m_b$ : body mass before infection.

